# Patterns of selection in the evolution of a transposable element

**DOI:** 10.1101/2021.09.06.459136

**Authors:** Julie Dazenière, Alexandros Bousios, Adam Eyre-Walker

**Author notes:** These authors contributed equally to this work (Julie Dazenière, Alexandros Bousios).

## Abstract

Transposable elements (TEs) are a major component of most eukaryotic genomes. Here, we present a new approach which allows us to study patterns of natural selection in the evolution of TEs over short time scales. The method uses the alignment of all elements with intact *gag/pol* genes of a TE family from a single genome. We predict that the ratio of non-synonymous to synonymous variants (vN/vS) in the alignment should decrease as a function of the frequency of the variants, because elements with non-synonymous variants that reduce transposition will have fewer progeny. We apply our method to Sirevirus LTR retrotransposons that are abundant in maize and other plant species and show that vN/vS declines as variant frequency increases, indicating that negative selection is acting strongly on the Sirevirus genome. The asymptotic value of vN/vS suggests that at least 85% of all non-synonymous mutations in the TE reduce transposition. Crucially, these patterns in vN/vS are only predicted to occur if the gene products from a particular TE insertion preferentially promote the transposition of the same insertion. Overall, this study is the first to use large numbers of intact elements to shed new light on the selective processes that act on TEs.

## Introduction

Transposable elements (TEs) are DNA sequences that can duplicate themselves and relocate from one chromosomal locus to another. They are divided into two main classes; class I elements (LTR and non-LTR retrotransposons) spread via a copy-and-paste pathway that involves an RNA intermediate, whereas class II elements (DNA transposons) transpose via a cut-and-paste pathway; both can result in a net increase of the TE copy number (1,2). TEs typically transmit vertically within hosts through the germline, but increasing evidence suggests that horizontal transfer of TEs can occur between species (3,4). Due to their activity over evolutionary time, TEs account for ∼50% of most primate genomes (5) and up to 80-90% of the genome of some plants (6,7). As such, TE activity is a major determinant of DNA sequence diversity and a key driver of species evolution (8,9).

TEs can be potentially harmful because they can integrate into genes and disrupt their function (10,11). They can also insert into promoters and regulatory sequences in the vicinity of genes, which can reduce expression levels by attracting silencing mechanisms and increasing local DNA methylation levels (12). TEs can also have negative consequences because they generate ectopic recombination (13,14), impose an energetic cost on genomes (15) and trigger intragenomic conflict when they capture fragments of host genes (16). Generally, TEs are thought to reduce host fitness and they have even been implicated with various diseases such as cancer in humans (10,17). However, TEs can potentially be advantageous for many of the same reasons, for example by providing exons or introducing promoter and enhancer elements near genes (18). In plants, there are several well-documented cases of agriculturally important traits that were caused by TE insertions (9,19–21).

TEs evolve in two separate dimensions. The first is through amplification within the host genome. After a new TE insertion occurs, it is polymorphic in the host population – some individuals have the element at this chromosomal position and others do not. This mutation, like all mutations, is subject to population processes of genetic drift and natural selection (13,22–24). The vast majority of these insertions will be lost from the population either because of genetic drift, or selection against them, if they are deleterious. A few insertions may also spread through the population, again either because of drift or selection, if the insertion is advantageous.

TEs also evolve in another dimension; they themselves evolve. This aspect of the evolutionary process has not been as extensively studied (4,25–29). It has been shown that the evolution of retrotransposons is largely dominated by negative selection both between and within families (4,25–29). Occasionally, positive adaptive evolution has been detected, as in the coiled coil region of ORFI of the human L1 LINE element (27). In contrast, there seems to be little evidence of selection acting on DNA transposons, except when these TEs are transferred between species (4).

When a new TE insertion occurs, it will start to accumulate mutations. These may be neutral with respect to transposition if the element inserts into a region of the genome from which it cannot further transpose, or if the changes have little effect on the probability of transposition; for example, if the mutations are synonymous. However, many mutations in the TE sequence will reduce the rate of transposition. As a consequence, as most TE insertions age so they will have fewer and fewer progeny; TEs are in a race to generate new copies of themselves before their sequence degenerates so that they can no longer transpose. All the TEs from a particular family in a single genome or in a population are connected to each other by a phylogeny. A consequence of the accumulation of mutations, which reduce transposition, is that internal branches in the tree should have fewer of these mutations, because internal branches represent elements that have successfully transposed (ignoring duplication of the locus), and internal branches with more daughter branches represent more successful elements. We can detect this pattern by considering the ratio of non-synonymous (vN) to synonymous (vS) variants, assuming that most synonymous mutations have no effect on transposition. We refer to mutations in the phylogeny as variants since they are neither substitutions nor polymorphisms; i.e. there is no guarantee that they are fixed in the species, as we would expect for a substitution, and although they are quite likely to be polymorphic within the population, because the element that the variant appears in is probably polymorphic in the population, the variant is defined relative to other copies of the element, and so referring to them as polymorphisms is inappropriate. We thus predict that branches internal to the tree should have lower vN/vS than external branches and that vN/vS should decline as a function of depth in the tree (i.e., branches with three descendant branches should have lower vN/vS on average than those with two). Occasionally a mutation might arise that increases the rate of transposition, and in this case the branch will have a high value of vN/vS.

An important caveat to these predictions for vN/vS is that the gene products of a TE insertion act in *cis* to generate copies of that particular locus, not copies of other loci of the same TE family in the genome. If the gene products from one insertion help in *trans* other TE loci to transpose, then even those with mutations that would render them otherwise incapable of transposition, will transpose and hence their vN/vS will be ∼1 (4,26).

It is well established that many TE copies contain debilitating mutations, such as stop codons and frameshifts in their coding sequences. What has rarely been demonstrated is the slow death of many TEs, through the accumulation of mutations that reduce transposition, and how those elements that avoid these, keep the lineage alive. In the only analysis of this kind Belshaw et al. (2004) showed that vN/vS was lower for the internal than external branches for human endogenous retroviruses (26).

We predict that vN/vS should typically be lower for internal than external branches. However, a challenge in the analysis of many TE families is their size and the speed at which they have expanded; inferring a robust phylogeny can therefore be difficult. We therefore developed a new method in which we align all the TE sequences from a single genome, and consider the variation in this alignment; in this alignment, we assume that a variant present in a single TE sequence appeared on a terminal branch of the tree (or it appeared on an internal branch and there was a back-mutation), one that is present in two copies occurred on a branch ancestral to two of the TEs in the genome…etc. Hence, we can infer the position at which the mutation occurred in the tree from its frequency. We therefore have two predictions. First, if synonymous mutations are neutral and non-synonymous mutations are neutral or deleterious in terms of TE transposition, then vN/vS should decline as a function of the frequency of the variant in the alignment. Second, if some mutations are advantageous to the TE, then this should lead to an increase in vN/vS amongst the highest frequency variants. Note that in contrast to an advantageous mutation spreading through a population, in which the advantageous allele can rapidly spread through the whole population, an advantageous mutation never spreads through all the TE copies in a single genome; the new variant can proliferate but there still remain all the elements that were already integrated into the genome. We test these predictions on a lineage of LTR retrotransposons that are found in plants. We focus only on the subset of elements that are potentially capable for autonomous transposition based on the completeness of their coding domain. We find clear evidence of negative selection, but no evidence of positive selection.

## Results

### TE datasets and plant species used in this study

We are interested in whether we can detect the signature of natural selection acting upon the sequence of transposable elements. We chose to investigate this among Sirevirus LTR retrotransposons, a TE lineage that is specific to plants (30) and often occupies a substantial proportion of the genome of their hosts (31). In the maize B73 genome, for example, there are five distinct Sirevirus families that collectively occupy ∼21% of the 2,300Mb genome (32). Among the five families, *Opie* and *Ji* have been very successful with each representing ∼10% of the genome (32) and with 10,778 and 10,563 full-length elements respectively (Table 1). In contrast, *Giepum, Hopie* and *Jienv* are found in much lower copy numbers (Table 1). These elements are considered full-length, because they contain all the structural features of complete LTR retrotransposons that are used by the various *de novo* TE identification pipelines: i.e. the presence of LTRs, a primer binding site, a polypurine tract, target-site duplication and the core domains of the *reverse transcriptase* or *integrase* genes (Supplementary Figure S1A). However, not all these elements are potentially functional due to mutations that can interrupt their genes. We consider that there is little point in testing for natural selection in elements that are clearly inactive based on their coding potential. We therefore identified a subset of elements that contain uninterrupted *gag* and *pol* open reading frames (see Methods) and refer to them as ‘intact’ (Supplementary Figure S1B). For *pol*, we further identified the junctions between *protease, integrase, reverse transcriptase* and *ribonuclease* so we could analyze them separately. In maize, this approach identified 2445, 2345, 140, 139 and 40 intact elements for *Opie, Ji, Hopie, Giepum* and *Jienv* respectively (Table 1). Because *Ji* and *Opie* are so much more numerous than *Hopie, Giepum* and *Jienv*, we randomly chose 130 sequences from *Opie* and *Ji* so as to have comparable population sizes across families for further analysis. Besides maize, we also identified full-length Sireviruses in a collection of monocot and eudicot species using the MASiVE annotation pipeline (33) and included in the analysis Sirevirus families that contained >5 intact elements (Table 1).

**Table 1.** Plant species and Sirevirus families included in this study. In maize, known TE exemplars were used to assign each element to a known family (see Methods). Simple names (e.g. Family 1, Family 2) were used for species with no exemplars.

| Species | Family | Full-length elements | Intact Sireviruses |
| --- | --- | --- | --- |
| *Zea mays* | *Opie* | 10788 | 2445 |
|  | *Ji* | 10563 | 2345 |
|  | *Hopie* | 374 | 140 |
|  | *Giepum* | 504 | 139 |
|  | *Jienv* | 279 | 40 |
| *Asparagus officinalis* | Family 1 | 92 | 22 |
|  | Family 2 | 457 | 58 |
| *Glycine max* | Family 1 | 403 | 64 |
|  | Family 2 | 842 | 404 |
| *Helianthus annuus* | Family 1 | 1360 | 60 |
| *Musa acuminata* | Family 1 | 263 | 134 |
| *Panicum hallii* var. *hallii* | Family 1 | 31 | 13 |
|  | Family 2 | 71 | 7 |
|  | Family 3 | 56 | 12 |
|  | Family 4 | 243 | 44 |
| *Sorghum bicolor* | Family 1 | 70 | 11 |
|  | Family 2 | 62 | 5 |
|  | Family 3 | 200 | 33 |
|  | Family 4 | 213 | 32 |

We aligned the genic sequences from each family in each species and then counted the number of synonymous and non-synonymous variants segregating at each codon site. Because we do not have a well resolved phylogenetic tree for the elements in each family, and hence we were unable to estimate rates of non-synonymous and synonymous change along each branch, we counted the number of synonymous and non-synonymous variants in groups of codons in which synonymous and non-synonymous variants were equally likely to occur and be scored (see Methods).

### Patterns of selection at the family level

We begin our analysis by considering patterns of evolution in the maize families and for all genes combined. Because high frequency variants are relatively rare, we combine frequency groups in the following scheme: we combine polymorphisms that have frequencies between 0 and 2^-6^, 2^-6^ and 2^-5^, 2^-5^ and 2^-4^ and so on until 2^-2^ and 2^-1^. In each family it is evident that vN/vS declines as a function of the frequency of the variants in the alignment across all five families before reaching an asymptote (Figure 1; Table 2). There is therefore a clear signature of negative selection acting in each Sirevirus family against non-synonymous variants. The value of vN/vS does not vary significantly between families for most frequency categories (Table 3).

**Table 2.** Spearman correlation between vN/vS and variant frequency for each family and each gene. Note the correlation is calculated across frequency categories.

| Group |  | Spearman's correlation coefficient | p-value |
| --- | --- | --- | --- |
| Families | Opie | -1 | 0.003 |
|  | Ji | -0.829 | 0.058 |
|  | Hopie | -0.6 | 0.208 |
|  | Giepum | -0.771 | 0.072 |
|  | Jienv | -0.7 | 0.188 |
| Genes | Gag | -0.829 | 0.058 |
|  | Integrase | -0.771 | 0.103 |
|  | Protease | -0.771 | 0.072 |
|  | Ribonuclease | -0.714 | 0.111 |
|  | Reverse transcriptase | -0.943 | 0.005 |

**Table 3.** Testing whether vN/vS differs between families and genes for each frequency category. The chi-square value is given along with the degrees of freedom and the p-value. Note, in the family analysis there are only 3 df in the lowest frequency class because *Jienv* has only 40 intact elements and hence no variants in the lowest frequency class.

| Analysis | Frequency category | Chi-square | df | p-value |
| --- | --- | --- | --- | --- |
| Between genes | $0 < x \leq 1/64$ | 12.09 | 4 | 0.02 |
| | $1/64 < x \leq 1/32$ | 8.69 | 4 | 0.07 |
| | $1/32 < x \leq 1/16$ | 10.65 | 4 | 0.03 |
| | $1/16 < x \leq 1/8$ | 15.2 | 4 | 0 |
| | $1/8 < x \leq 1/4$ | 9.89 | 4 | 0.04 |
| | $1/4 < x \leq 1/2$ | 11.09 | 4 | 0.03 |
|  | Total | 67.61 | 24 | < .00001 |
| Between families | $0 < x \leq 1/64$ | 7.99 | 3 | 0.05 |
| | $1/64 < x \leq 1/32$ | 7.06 | 4 | 0.13 |
| | $1/32 < x \leq 1/16$ | 9.52 | 4 | 0.05 |
| | $1/16 < x \leq 1/8$ | 5.9 | 4 | 0.21 |
| | $1/8 < x \leq 1/4$ | 3.29 | 4 | 0.51 |
| | $1/4 < x \leq 1/2$ | 2.91 | 4 | 0.57 |
|  | Total | 36.67 | 23 | 0.035 |

**Figure 1.**
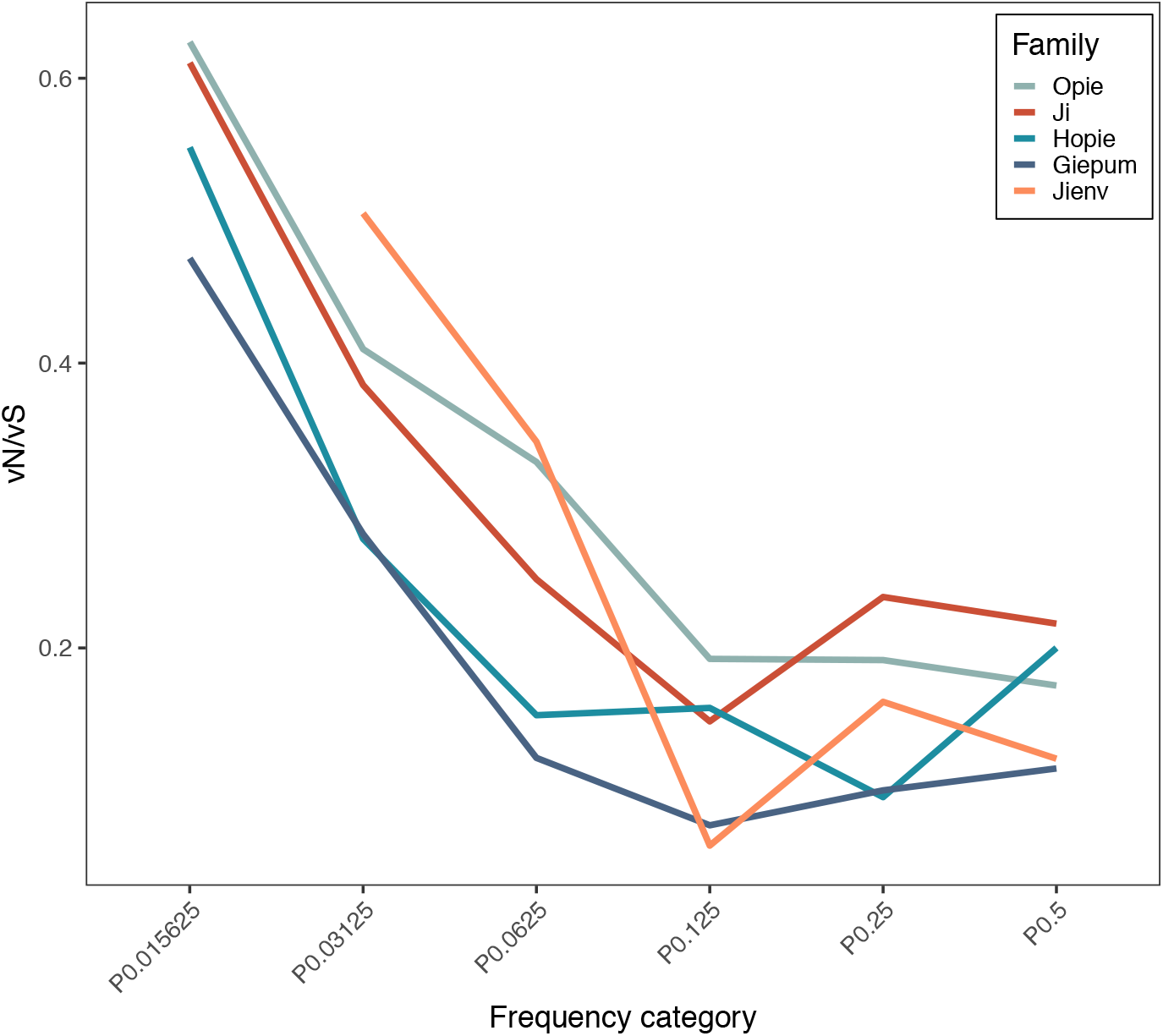
The value of vN/vS as a function of the frequency of the variants in the alignment for the five families of Sireviruses in maize is shown with plain lines. *Ji* and *Opie* are sampled to 130 sequences each, while *Giepum, Hopie* and *Jienv* are the full datasets.

The patterns of vN/vS are remarkably similar in the different families, even though two of the families are much more numerous than the others. A factor that might influence the patterns we observe is the age of the elements. We can potentially estimate the relative age of each element from the divergence between the two LTRs that flank each element; it is assumed that these are identical when the element first inserts and hence divergence between the LTRs can be used to estimate the relative age of each element; note that we do not attempt to estimate the absolute age because the LTRs might have a function and hence evolve more slowly than the mutation rate. We find significant differences in the median relative age between families (Kruskal-Wallis test, p<0.001) (Figure 2), with pairwise Mann-Whitney tests suggesting that the median age of *Opie* elements is significantly different to *Ji* (p<0.001), and to *Hopie* (p<0.001). However, the differences in age are quite small.

**Figure 2.**
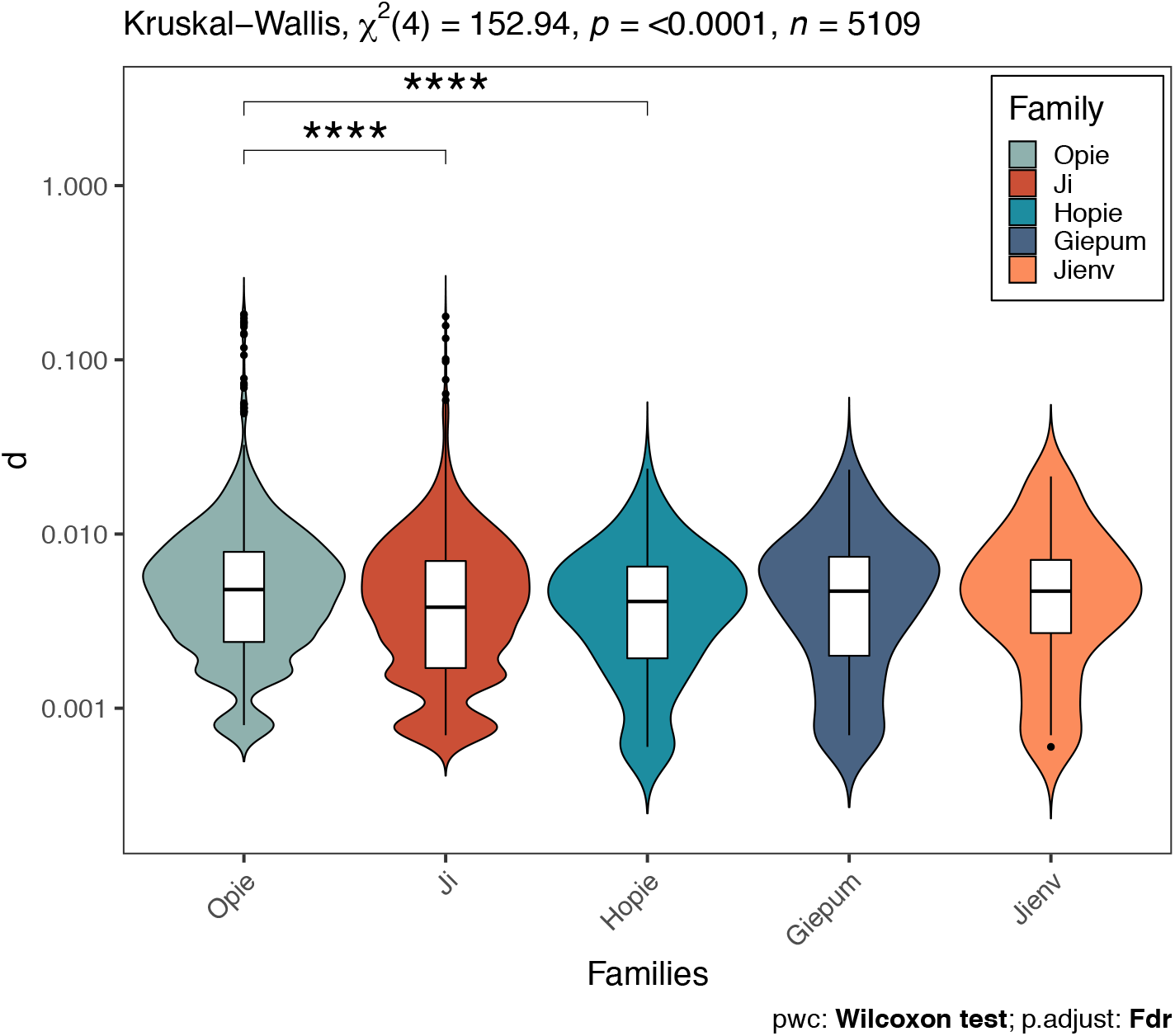
The distribution of relative ages across TE copies from each family. Violin plots show the distribution of divergence (numbers of point mutations and indels per site) between the LTRs of each element. Inset is a boxplot showing the medians and inter-quartile range (IQR). The whiskers go from the hinge to the largest/smallest value no further than 1.5*IQR. Note the log-scale of the Y-axis.

### Patterns of selection acting on the TE genes

We now turn attention to whether there are differences in the pattern of natural selection between the genes in the Sirevirus sequence, by summing data across families for each gene. As we might have expected from the family analysis, we find that vN/vS declines significantly over the first four frequency categories before increasing slightly in four of the genes - *gag, integrase, protease* and *ribonuclease* (Figure 3A; Table 2). This is the signal we might expect from adaptive evolution, however, these increases are likely due to sampling error because the number of non-synonymous variants involved in each case is small - 50, 12, 1 and 3 non-synonymous variants have frequencies in excess ⅛ in the *gag, integrase, protease* and *ribonuclease* genes respectively. The value of vN/vS varies significantly between the genes for all frequency categories (Table 3) with *integrase* and especially *gag* being less conserved than the other three genes. For *gag*, this is also reflected in the much higher length variation among families compared to the other genes (Figure 3B). The average value of vN/vS over the last three frequency categories is 0.15 for the five genes.

**Figure 3.**
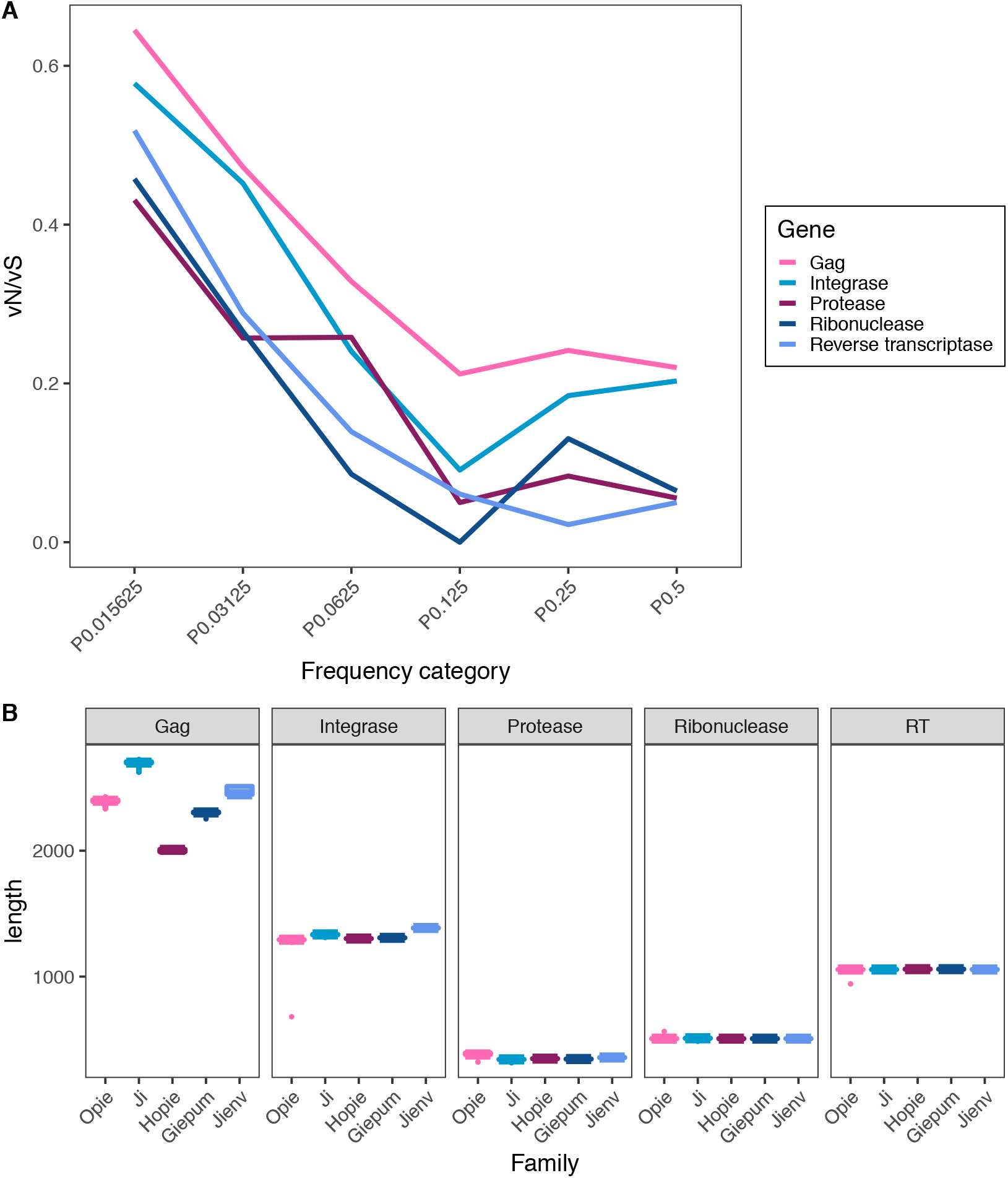
A) The value of vN/vS as a function of the frequency of the variants in the alignment for the 5 genes in the Sirevirus element, with the families combined. B) The length variation of the 5 maize families for each gene.

### Patterns of selection in other plant hosts

It is of interest to see if these general patterns are found in other species. We extracted Sireviruses from six other species which have between 31 and 1360 full-length elements and between 5 and 398 intact elements (Table 1). As in maize we find that vN/vS is highest in the singleton class and then declines across frequency classes before coming to an asymptote (Figure 4).

**Figure 4.**
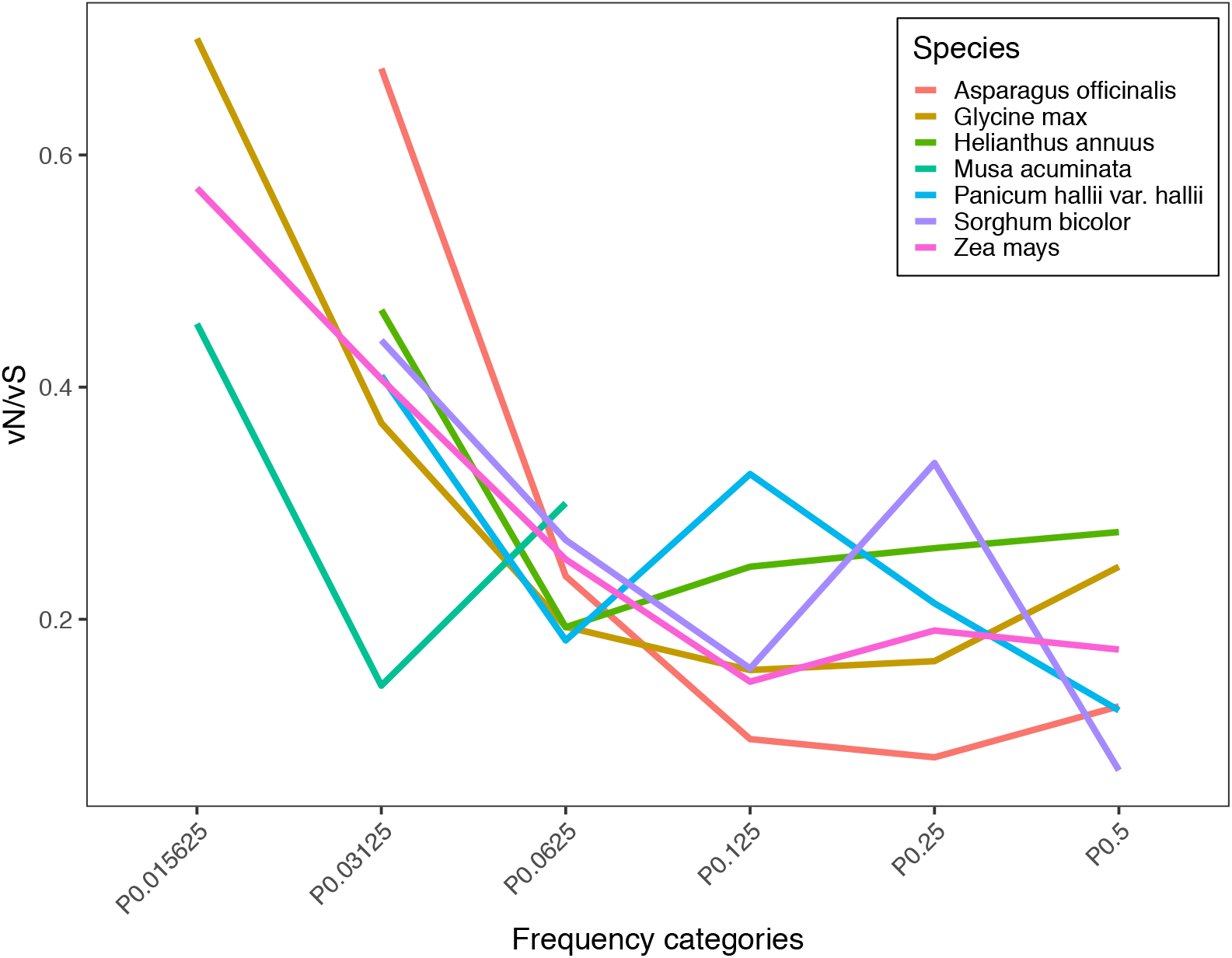
The value of vN/vS as a function of the frequency of the variants in the alignment for Sirevirus families in various plant species.

## Discussion

We have investigated patterns of selection in intact Sirevirus elements within the genome of maize and other species using a new approach. We have aligned TE sequences from a single genome and considered the ratio of non-synonymous to synonymous variants as a function of the frequency of the variants in the alignment. We find that the ratio of non-synonymous to synonymous variants, vN/vS, declines as a function of the frequency of the variant in an alignment of the TE sequences in a single genome. This is expected; TE sequences that tend to accumulate deleterious non-synonymous variants are likely to be less able to transpose. The value of vN/vS across all genes for variants with frequencies in excess of 1/8 is 0.15 and this implies that at least 85% of non-synonymous mutations in the Sirevirus sequence reduces transposition. The reason is as follows; let us assume that synonymous mutations are neutral (i.e. they have no effect on the rate of transposition), then the rate at which synonymous mutations accumulate is proportional to the mutation rate, *u*. In contrast, let us assume that non-synonymous mutations are neutral or deleterious, in the sense that they reduce transposition; let the proportion that are neutral be *f* then the rate at which non-synonymous variants accumulate is proportional to *uf* and hence the ratio of vN/vS is simply an estimate of *f*; hence if vN/vS = *f* = 0.15, this implies that 15% are neutral and 85% are deleterious. This is likely to be a lower estimate because some non-synonymous mutations might increase the rate of transposition and some synonymous mutations might decrease the rate.

One of the challenges for any TE is avoiding parasitism. A functional and active TE will produce gene products that will allow it to generate new copies of itself that can be integrated into the host genome at new locations. However, these gene products can potentially be used by other elements to transpose themselves; a number of very successful TEs, such as SINEs in mammals and MITEs in plants, are incapable of transposing themselves, and duplicate by parasitising the machinery of other autonomous TEs - LINEs and MLEs in the case of the SINEs (34) and MITEs (35) respectively. It is clearly in the interest of a TE to avoid this parasitism; the more gene products get diverted to other parasitic elements, the less likely the element that produced the gene products is to successfully transpose itself. The observation that vN/vS is very substantially less than one amongst the high frequency variants suggests that each Sirevirus copy is successful at targeting most of its own gene products to their own transposition (26). The reason is as follows. If the gene products from a particular TE diffused throughout the cell, and helped other TE copies to transpose, then this would allow TEs to transpose that did not produce any useful gene product including those with substantial numbers of non-synonymous mutations. Hence vN/vS would be substantially higher and we would not observe vN/vS declining as a function of the frequency of the variants in the alignment. Hence our observation that vN/vS<1 for higher frequency categories is evidence for this *cis*-targeting as Belshaw et al. (2004) first noted. This cis-targeting has been experimentally confirmed for LINE elements (36,37), but there is no evidence for this *cis*-targeting for the one LTR retrotransposon family that has been investigated in depth, the Ty1 in yeast (38). In addition, the life cycle of an LTR retrotransposon makes it difficult to see how *cis*-targeting could be brought about (39). Zhang et al. (2020) have hypothesized that *cis* preference might arise if just one or two elements are transcribed in the germ-line, even if a given family has numerous copies in the genome. This hypothesis requires further testing, but it is supported from preliminary evidence of a recent study on Arabidopsis and maize TEs (40). In this study, the authors mapped long-read libraries of full-length mRNAs to TEs in an effort to pinpoint which copies of a family are truly expressed. This is not possible using short read RNA-seq data due to the multimapping effect on TEs (41). The authors found that only 4% of all annotated TEs in arabidopsis were expressed in a triple mutant that removes many layers of epigenetic silencing. In maize, they focused on *Opie*, and using libraries from different tissues they found that only 6 copies were expressed out of a total of ∼12,000. It is noteworthy that, similar to LTR retrotransposons, DNA transposons cannot perform *cis* targeting because of their life cycle – the transposase is produced in the cytoplasm and diffuses back into the nucleus to cut and paste the element – and DNA transposons show vN/vS = 1 within a species (4).

Our method has the potential to detect periods of adaptive evolution. If a TE undergoes a non-synonymous mutation which allows the TE to transpose more often or which allows the TE to survive, then this TE will have more progeny. Such mutations are more likely to be non-synonymous and hence we might expect to see an elevation in vN/vS amongst high frequency variants. There are, however, two problems. First, negative selection is expected to lead to a decrease in vN/vS across frequency categories, so this may mask the signature of positive selection. Second, different advantageous mutations will have different effects; for example, one might lead to an increase in transposition such that 40% of the elements carry the advantageous mutation, whereas another might lead to only 10% of the TE population carrying the mutation; i.e. the signature of adaptive evolution is likely to be spread across many frequency categories.

The method makes a number of simplifying assumptions. We assume that the only manner in which a TE can make a copy of itself is through transposition, rather than through duplication of the genome, chromosome or part of the chromosome. We also assume that there is little or no gene conversion between TEs. Making these assumptions is unlikely to affect our results; both processes will tend to introduce noise in the analysis; i.e. we might have a TE which is incapable of transposition and which is accumulating non-synonymous and synonymous mutations at equal rates; all mutations should appear as singletons, unless there is duplication or gene conversion, which can potentially change a singleton into a two copies, hence elevating vN/vS in higher frequency categories.

It is conspicuous that the age distributions of the three smaller and the larger two Sirevirus families are remarkably similar. This is unexpected because one would expect these families to be transposing independently, and our vN/vS results support the *cis* action of the TE protein products. What then could generate the similarity in the age profiles? There are two possibilities. The host is suppressing families in concert, and second the age profiles represent an equilibrium state in which the rate of transposition and deletion of elements has been constant for some time. This however, is itself surprising because under this hypothesis we would expect most elements to be young because most mutations are young in a population, and these are lost rapidly by genetic drift or selection. However, in most studies that report the age distribution of LTR retrotransposon families we see that this is not the case; there is a dearth of elements that are very young. This might arise because the LTRs are not always perfect copies when they are first formed.

We have shown that the number of non-synonymous versus synonymous variants in an alignment of TE sequences from a single genome, declines as a function of the frequency of the variants in the alignment. This is consistent with the action of negative selection; elements that accumulate non-synonymous mutations are less likely to transpose and hence have progeny, and hence have a high frequency in the alignment. This is the first demonstration of how natural selection acts upon an element during its evolution over short timescales.

## Materials and Methods

### Identification of intact Sirevirus elements

We run MASiVE (33) to identify full-length Sirevirus elements in the genomes of the seven species used in this study. Supplementary Table S4 contains information on the genome versions, source links and citations for these species. Similar to other *de novo* LTR retrotransposon identification algorithms, MASiVE identifies full-length Sireviruses based on the presence of structural features (e.g. LTRs, primer binding site, polypurine tract, target-site duplication) and positive hits with the core domain of the *reverse transcriptase* and *integrase* genes using the Pfam Hidden Markov Models (HMM) PF07727.9 and PF00665.21. However, it is not guaranteed that these elements are intact in terms of their coding domains and if they are competent for autonomous transposition (Figure S1; in fact, it is likely that most elements have acquired one or more mutations after integration in the genome that disrupted the *gag* and *pol* ORFs). Generally, the proportion of TEs within a family that are intact elements is unknown and identifying them requires substantial resources and TE expertise.

For this study it was necessary to characterize these elements and, hence, we devised the following pipeline: For every species, we first produced a multiple alignment using Mafft G-INS-i algorithm (42) of all Sirevirus elements based on the HMM domain of the *reverse transcriptase* gene. Due to the high numbers of elements in maize (Table 1), we used the CD-HIT clustering package (43) to reduce the number of elements prior to the alignment. We required a 95% identity threshold (-c 0.95) and a coverage of at least 90% for every sequence pair (-aL and -aS both at 0.9). Every sequence was placed in the most similar cluster and not the first one that meet the thresholds (–g 1). We then run FastTree (44) to generate maximum likelihood phylogenetic trees, which were visualized using FigTree (http://tree.bio.ed.ac.uk/software/figtree/, version 1.4.3). We assigned elements into families based on the branching pattern and bootstrap support. The addition of known Sirevirus exemplars in maize from (45) and the maize TE database was used to assign branches to specific family names. We then run getorf from the EMBOSS suite (46) with -minsize 1000 to identify long open reading frames (ORFs) within the internal domain (i.e. excluding the LTRs) of each element and hmmscan from the HMMER software (hmmer.org) using a list of known HMMs for LTR retrotransposons (Table S5) to annotate the ORFs as part of the *gag* or *pol* polyprotein. The length and start positions of the *gag* and *pol* ORFs were then plotted, while for *pol* we additionally required for the presence of the amino acid motif ADIFTK that is conserved among *Copia* LTR retrotransposons (47). The motif lies a short distance upstream of the 3’ end of the *pol* gene and was therefore used as an anchor to only keep *pol* ORFs that were complete on the 3’ end. An example of this process is shown in Supplementary Figure S2.

Finally, the junctions of the four genes within *pol* were identified as follows: *protease* was from the beginning of *pol* up till the beginning of the GAG-pre-integrase PFAM domain (PF13976), which was hence defined as the *protease*/*integrase* junction. This matches the boundaries of these two genes as identified by Peterson-Burch et al. (2002) (48). The C-terminus of the *integrase* is generally poorly conserved across *Copia* elements, but Sireviruses contain the ILGD motif a short distance (10-20 amino-acids) upstream of the 3’ end of the gene (48). In maize Sireviruses, this motif is followed after ∼50 amino-acids by the beginning of the *reverse transcriptase* PFAM domain (PF07727). We therefore approximately set the *integrase*/*reverse transcriptase* junction to be 30 amino-acids upstream of PF07727. The junction of the *reverse transcriptase* with the *ribonuclease* is also not precisely defined in the literature. However, *ribonuclease* starts with a highly conserved D_10_E_48_D_70_ motif (49), and the region of the first aspartic acid can be readily identified in Sireviruses. The aspartic acid also lies ∼85 amino-acids downstream of the *reverse transcriptase* PFAM domain (PF07727) and it does not overlap the last conserved domain of the *reverse transcriptase* gene as it was identified by (50). We approximately set the *reverse transcriptase* with the *ribonuclease* junction to be 15 amino-acids upstream of the first aspartic acid of the D_10_E_48_D_70_ motif.

### Multiple sequences alignments

Our method relies on the number of non-synonymous and synonymous sites identified in a multiple sequence alignment (MSA). The TE sequences in each family are moderately divergent from each other and they contain a number of indel mutations, so aligning them was challenging. We tried several approaches; MAFFT (42) and MACSE (51) both introduced substantial numbers of gaps that were not multiples of three. We therefore used TranslatorX (52) in association with MUSCLE (53) to align the sequences at the amino acid level. Visual inspection suggested that these alignments were reasonable; i.e. by aligning at the amino acid level we do not allow indels that introduce a frameshift, but frameshifts are apparent as sections of the alignment which align poorly in the sequences that are genuinely frameshifted. We found only one such case in the *gag* gene of the *Opie* family. Such alignment problems will introduce noise into our analysis, by generating non-synonymous and synonymous variants at the same frequency.

Our alignments contain multiple gaps in certain regions. To investigate whether the quality of our alignments affected our results we repeated our analysis. First, our intact elements vary in length, and so we chose only those sequences within our selected range (Figure S2) whose length class contained at least 10 sequences. We then realigned the sequences. Second, we edited the original alignment to remove those sections that had multiple gaps. Our results remained qualitatively unchanged, so we proceeded with the original alignments.

### Determination of the vN/vS ratio

We want to estimate the rate at which non-synonymous and synonymous variants accumulate in the TE sequences. One option would be to construct the phylogeny of the TE sequences and then estimate the rate at which variants accumulate using one of the many methods which have been developed to estimate rates of synonymous and non-synonymous substitution. However, the phylogeny is poorly resolved for our TE families since they are relatively young. We therefore developed a simple counting method in which synonymous and non-synonymous mutations were equally likely to be appear and be counted. To do this we focussed on groups of codons, generally four codons. For example, the four codons from Phenylalanine and Leucine - TTT, TTC, CTT and CTC. Here the synonymous and non-synonymous mutations involve the same mutation C<>T, and hence are expected to have the same mutation rate (ignoring context effects) and we can score both synonymous and non-synonymous variants even if they occur together (Table 5). We had five sets of four codons in which the synonymous and non-synonymous mutations were the same, and three sets of four codons in which the synonymous and non-synonymous mutations were the complement of each other; for example, the Isoleucine and Valine codons ATT, ATC, GTT and GTC; here the synonymous mutation is T<>C and the non-synonymous mutation is it’s complement A<>G. We also included one set of 16 codons, the codons of Proline, Threonine, Alanine and 4-fold degenerate codons of Serine (i.e. all codons of the form NCN). Here synonymous and non-synonymous mutations are expected to occur at equal rates (assuming context effects are minimal); for example, a TCT codon is equally likely to give rise to TCA and ACT, representing synonymous and non-synonymous changes. The list of codons is given in Supplementary Table S3. In some cases, the group of 16 codons could give rise to tri-allelic sites. In these cases we took the frequency of the rarest allele. All of these sets are independent of each other - they do not share any codons in common. There are other sets that we could use but unfortunately these are not independent. For a codon site to be included in the analysis it had to contain at least 10 instances of a set of codons (see codon 4 in Table 5). However, one codon site could contribute to multiple codon sets (see codon 5 in Table 5). In terms of the frequency of the variant we always consider the minor allele, and the frequency is considered across the whole alignment, not just the set of codons in which the variant appears. For example, codon 5 in Table 5 has a synonymous variant at a frequency of 1 in 10; let us imagine that there are 100 sequences; the variant frequency would then be 1 in 100.

**Table 5.** Examples of how synonymous and nonsynonymous variants are counted.

| Sequence | Codon 1 | Codon 2 | Codon 3 | Codon 4 | Codon 5 |
| --- | --- | --- | --- | --- | --- |
| 1 | TTT | TTT | TTT | TTT | TTT |
| 2 | TTT | TTT | TTC | TTC | TTC |
| 3 | TTC | CTT | CTT | AGA | CTT |
| ... | ... | ... | ... | ... | ... |
| 10 | TTC | CTT | CTT | AGG | CTT |
| 11 | TTC | CTT | CTT | GCT | AAA |
| 12 | TTC | CTT | CTT | ACT | AAA |
| ... | ... | ... | ... | ... |  |
| 20 | TTC | CTT | CTT | TTT | AAG |
| Synonymous variant count | 1 | 0 | 1 | 0 | 2 |
| Non-synonymous variant count | 0 | 1 | 1 | 0 | 1 |
| Notes |  |  |  | No codon set has 10 instances | Multiple codon sets included |

### Sampling of Opie and Ji

The sampling of the two biggest families of maize, *Opie* and *Ji*, was performed on the dataset of intact elements. For each of these two families, we sampled of 130 sequences at random. The pipeline was then applied as previously described.

## Supplementary data

**Supplementary Figure S1.**
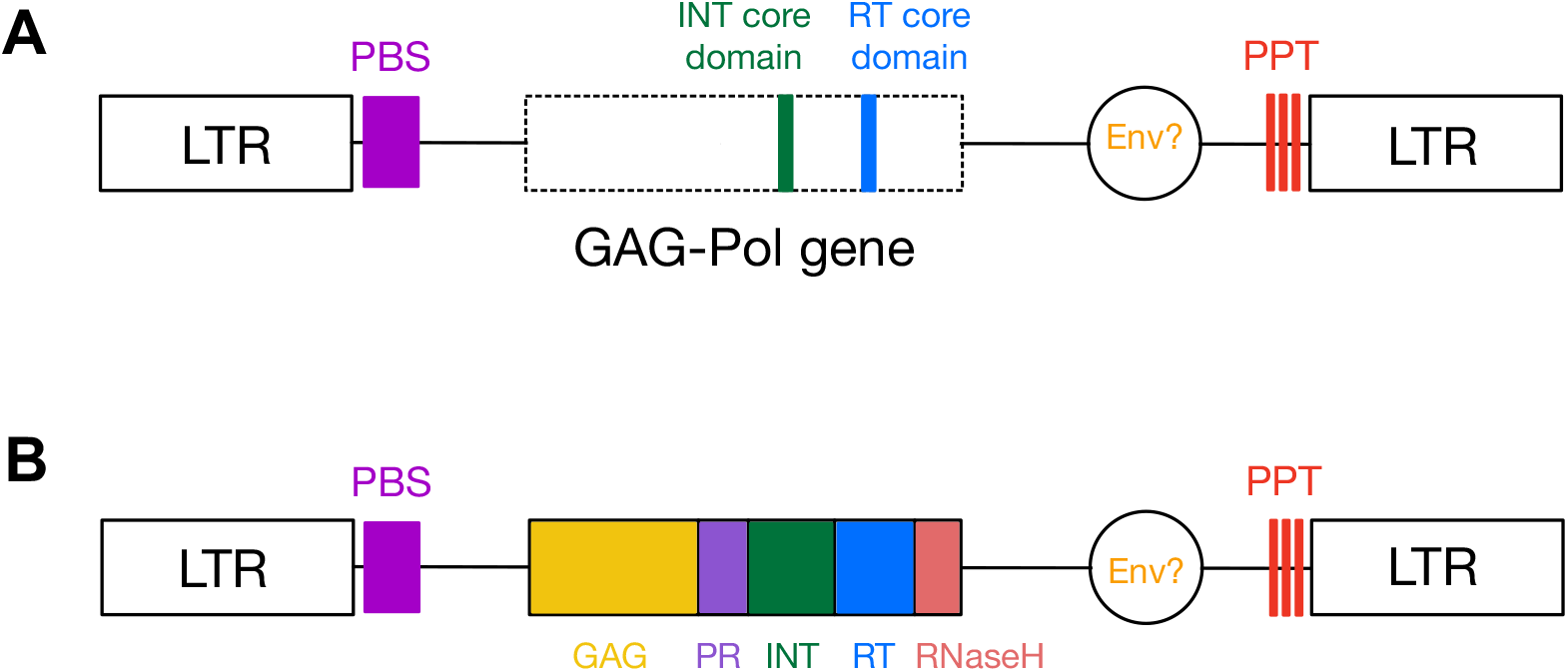
Difference between a A) full-length Sirevirus LTR-retrotransposons and B) an intact element as defined in this work. The figures show the relative positions of some key features of a Sirevirus TE ; the primer binding site (PBS), the polypurine tract (PPT), the *gag* gene, and the *pol* gene, composed of 4 genes: the *protease* (PR), *integrase* (INT), *reverse transcriptase* (RT) and *ribonuclease-H* (RNaseH) genes. Note that some Sireviruses possess an envelope-like gene (Env) downstream the *gag-pol* domain.

**Supplementary Figure S2.**
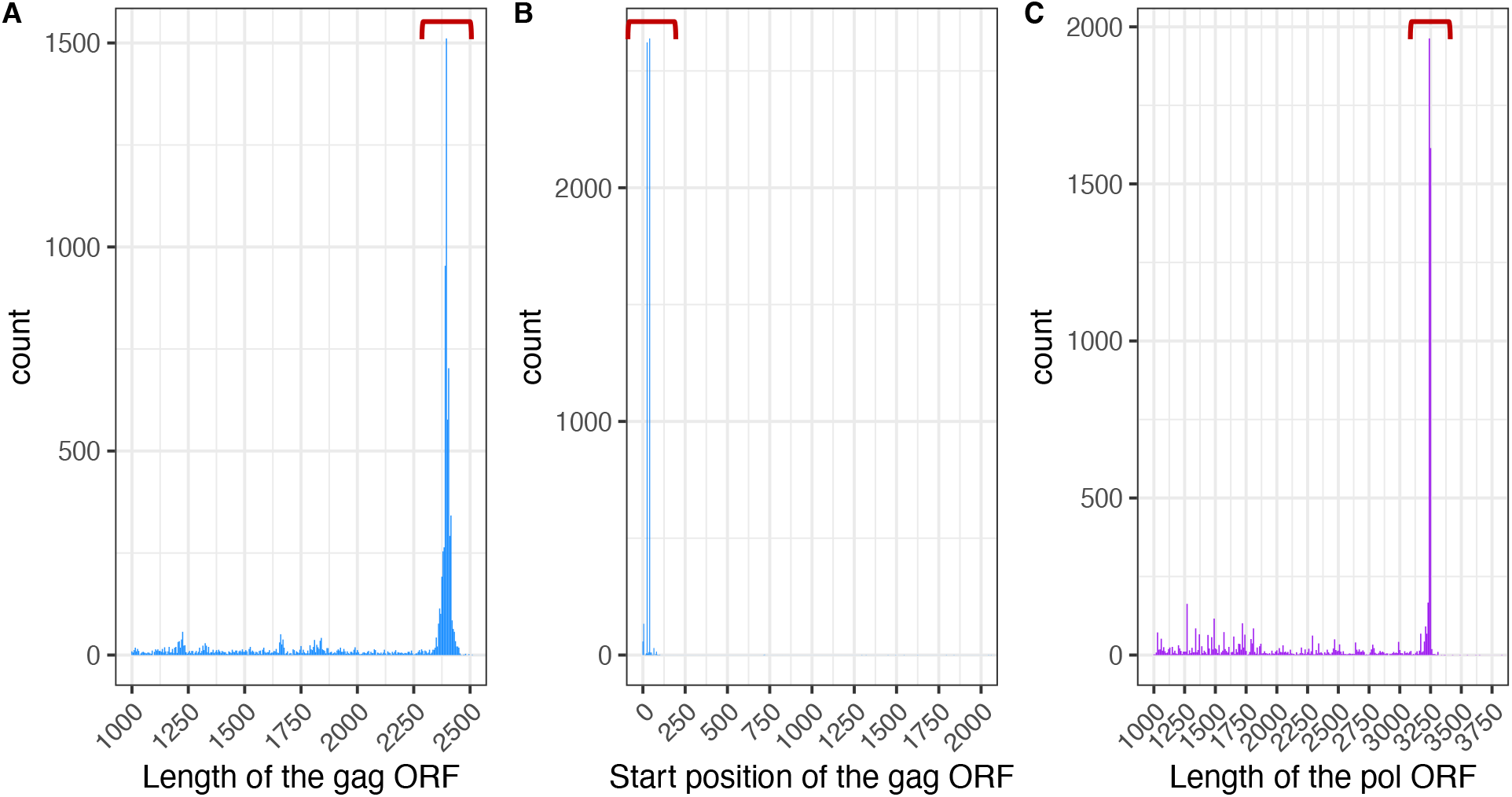
Method to retrieve the intact ORF for *gag* (A and B) and *pol* (C) in the *Opie* family. ORFs were first categorized as *gag* and *pol* based on their overlap with TE-specific HMM motifs (shown in Table S5). See Methods for more details about the pipeline. (A) shows the selected length for the ORF of the *gag* gene (highlighted with the red bracket), (B) which were further selected and plotted to visualize their starting positions. The chosen starting position of the complete *gag* ORFs are highlighted with the red bracket. (C) In *pol*, we only select a specific length and not a starting position of the ORF, because the starting position within the internal domain can vary substantially within a family given individual insertions and deletions that elements may have acquired after integration. All *pol* ORFs contain the amino acid motif ADIFTK that is conserved among *Copia* LTR retrotransposons.

**Supplementary Table 1.** Binomial test for each family, the number of NS and S sites for the genes were combined within each family.

| Family | Frequency category | Non-synonymous sites | Synonymous sites | p-value |
| --- | --- | --- | --- | --- |
| Giepum | $0 < x \leq 1/64$ | 215 | 454 | 0 |
| | $1/64 < x \leq 1/32$ | 21 | 75 | 0 |
| | $1/32 < x \leq 1/16$ | 7 | 57 | 0 |
| | $1/16 < x \leq 1/8$ | 4 | 53 | 0 |
| | $1/8 < x \leq 1/4$ | 3 | 30 | 0 |
| | $1/4 < x \leq 1/2$ | 3 | 26 | 0 |
| Hopie | $0 < x \leq 1/64$ | 209 | 379 | 0 |
| | $1/64 < x \leq 1/32$ | 13 | 47 | 0 |
| | $1/32 < x \leq 1/16$ | 11 | 72 | 0 |
| | $1/16 < x \leq 1/8$ | 6 | 38 | 0 |
| | $1/8 < x \leq 1/4$ | 2 | 21 | 0 |
| | $1/4 < x \leq 1/2$ | 5 | 25 | 0 |
| Jienv | $0 < x \leq 1/64$ | 0 | 0 | 1 |
| | $1/64 < x \leq 1/32$ | 98 | 194 | 0 |
| | $1/32 < x \leq 1/16$ | 20 | 58 | 0 |
| | $1/16 < x \leq 1/8$ | 3 | 49 | 0 |
| | $1/8 < x \leq 1/4$ | 6 | 37 | 0 |
| | $1/4 < x \leq 1/2$ | 11 | 90 | 0 |
| Opie | $0 < x \leq 1/64$ | 324 | 518 | 0 |
| | $1/64 < x \leq 1/32$ | 50 | 122 | 0 |
| | $1/32 < x \leq 1/16$ | 40 | 121 | 0 |
| | $1/16 < x \leq 1/8$ | 30 | 156 | 0 |
| | $1/8 < x \leq 1/4$ | 27 | 141 | 0 |
| | $1/4 < x \leq 1/2$ | 25 | 144 | 0 |
| Ji | $0 < x \leq 1/64$ | 372 | 609 | 0 |
| | $1/64 < x \leq 1/32$ | 65 | 169 | 0 |
| | $1/32 < x \leq 1/16$ | 36 | 145 | 0 |
| | $1/16 < x \leq 1/8$ | 19 | 128 | 0 |
| | $1/8 < x \leq 1/4$ | 29 | 123 | 0 |
| | $1/4 < x \leq 1/2$ | 28 | 129 | 0 |

**Supplementary Table 2.** Chi-square test between Hopie and Giepum and a random replicate of Ji and Opie downsampled.

| Analysis | Frequency category | Chi-square | df | p-value |
| --- | --- | --- | --- | --- |
| giepum/ji | $0 < x \leq 1/64$ | 5.55 | 1 | 0.02 |
| | $1/64 < x \leq 1/32$ | 0.94 | 1 | 0.33 |
| | $1/32 < x \leq 1/16$ | 2.04 | 1 | 0.15 |
| | $1/16 < x \leq 1/8$ | 0.9 | 1 | 0.34 |
| | $1/8 < x \leq 1/4$ | 1.26 | 1 | 0.26 |
| | $1/4 < x \leq 1/2$ | 0.52 | 1 | 0.47 |
|  | Total | 11.21 | 6 | 0.082 |
| giepum/opie | $0 < x \leq 1/64$ | 6.26 | 1 | 0.01 |
| | $1/64 < x \leq 1/32$ | 1.29 | 1 | 0.26 |
| | $1/32 < x \leq 1/16$ | 4.55 | 1 | 0.03 |
| | $1/16 < x \leq 1/8$ | 2.3 | 1 | 0.13 |
| | $1/8 < x \leq 1/4$ | 0.58 | 1 | 0.45 |
| | $1/4 < x \leq 1/2$ | 0.12 | 1 | 0.73 |
|  | Total | 15.1 | 6 | 0.019 |
| hopie/ji | $0 < x \leq 1/64$ | 0.79 | 1 | 0.37 |
| | $1/64 < x \leq 1/32$ | 0.63 | 1 | 0.43 |
| | $1/32 < x \leq 1/16$ | 1.29 | 1 | 0.26 |
| | $1/16 < x \leq 1/8$ | 0 | 1 | 1 |
| | $1/8 < x \leq 1/4$ | 0.85 | 1 | 0.36 |
| | $1/4 < x \leq 1/2$ | 0 | 1 | 1 |
|  | Total | 3.56 | 6 | 0.736 |
| hopie/opie | $0 < x \leq 1/64$ | 1.15 | 1 | 0.28 |
| | $1/64 < x \leq 1/32$ | 0.89 | 1 | 0.35 |
| | $1/32 < x \leq 1/16$ | 3.78 | 1 | 0.05 |
| | $1/16 < x \leq 1/8$ | 0.03 | 1 | 0.86 |
| | $1/8 < x \leq 1/4$ | 0.38 | 1 | 0.54 |
| | $1/4 < x \leq 1/2$ | 0 | 1 | 1 |
|  | Total | 6.23 | 6 | 0.398 |

**Supplementary Table 3.** Codon sets used to count the numbers of synonymous and non-synonymous variants.

| Codon set | Amino acids | Number of codons | Codons |
| --- | --- | --- | --- |
| 1 | Ser/Pro/Thr/Ala | 16 | TCT, TCC, TCA, TCG, CCT, CCC, CCA, CCG, ACT, ACC, ACA, ACG, GCT, GCC, GCA, GCG |
| 2 | Phe, Leu | 4 | TTT, TTC, CTT, CTC |
| 3 | Tyr/His | 4 | TAT, TAC, CAT, CAC |
| 4 | Lys/Glu | 4 | AAA, AAG, GAA, GAG |
| 5 | Cys/Arg | 4 | TGT, TGC, CGT, CGC |
| 6 | Arg/Gly | 4 | AGA, AGG, GGA, GGG |
| 7 | Iso/Val | 4 | ATT, ATC, GTT, GTC |
| 8 | Asn/Asp | 4 | AAT, AAC, GAT, GAC |
| 9 | Ser/Gly | 4 | AGT, AGC, GGT, GGC |

**Supplementary Table 4.** List of the genomes used for this analysis.

| Species | Website (accession date) | Publication (PubMed ID) |
| --- | --- | --- |
| Zea mays B73 | http://ensembl.gramene.org/Zea_mays/Info/Index (June 2018) | 19965430 |
| Musa acuminata | http://banana-genome-hub.southgreen.fr/download (May 2016) | 26984673 |
| Glycine max | http://genome.jgi.doe.gov/pages/dynamicOrganismDownload.jsf?organism=Gmax (May 2016) | 20075913 |
| Asparagus officinalis | https://genome.jgi.doe.gov/portal/pages/dynamicOrganismDownload.jsf?organism=Aofficinalis (June 2018) | 29093472 |
| Helianthus annuus | https://genome.jgi.doe.gov/portal/pages/dynamicOrganismDownload.jsf?organism=Hannuus (June 2018) | 28538728 |
| Panicum hallii var. Hallii | https://genome.jgi.doe.gov/portal/pages/dynamicOrganismDownload.jsf?organism=PhalliiHAL (June 2018) | 30523281 |
| Sorghum bicolor | https://genome.jgi.doe.gov/portal/pages/dynamicOrganismDownload.jsf?organism=Sbicolor (June 2018) | 29161754 |

**Supplementary table 5.** List of the Pfam HMM models used to determine the ORFs of each gene.

| Pfam ID | Description |
| --- | --- |
| PF14223 | *gag*-polypeptide of LTR copia-type |
| PF13961 | Domain of unknown function |
| PF00098 | Zinc finger |
| PF07727 | Reverse transcriptase (RNA-dependent DNA polymerase) |
| PF00665 | Integrase core domain |
| PF13976 | GAG-pre-integrase domain |

